# Incorporation of Epstein-Barr viral variation implicates significance of LMP1 in survival prediction and prognostic subgrouping in Burkitt lymphoma

**DOI:** 10.1101/2024.05.03.592343

**Authors:** Isaac E. Kim, Cliff Oduor, Julian Stamp, Micah A. Luftig, Ann M. Moormann, Lorin Crawford, Jeffrey A. Bailey

## Abstract

While Epstein-Barr virus (EBV) plays a role in Burkitt lymphoma (BL) tumorigenesis, it is unclear if EBV genetic variation impacts clinical outcomes. From 130 publicly available whole-genome tumor sequences of EBV-positive BL patients, we used least absolute shrinkage and selection operator (LASSO) regression and Bayesian variable selection models within a Cox proportional hazards framework to select the top EBV variants, putative driver genes, and clinical features associated with patient survival time. These features were incorporated into survival prediction and prognostic subgrouping models. Our model yielded 22 EBV variants including seven in LMP1 as most associated with patient survival time. Using the top EBV variants, driver genes, and clinical features, we defined three prognostic subgroups that demonstrated differential survival rates, laying the foundation for incorporating EBV variants such as those in LMP1 as predictive biomarker candidates in future studies.

## INTRODUCTION

Burkitt lymphoma (BL) comprises 30-50% of all pediatric cancers in equatorial Africa.^1,2^ Molecularly characterized by a translocation of the immunoglobulin (Ig) enhancer upstream of the MYC gene, BL has traditionally been categorized into three epidemiologic subtypes (endemic BL, sporadic BL, and HIV-BL).^3^ Sporadic BL (sBL) is most often diagnosed in adolescents, adults, and elderly patients residing in non-tropical countries and has a survival rate greater than 90%, unlike eBL, which is primarily diagnosed in children under nine years of age living in equatorial Africa and has a lower survival rate of 50%, likely largely due to access to care.^4^ It has been hypothesized that repeated infections with the malarial parasite, Plasmodium falciparum, combined with early-age infection with Epstein-Barr virus (EBV), induces polyclonal B-cell expansion and activation-induced cytidine deaminase (AID)-dependent DNA damage, increasing the probability of the MYC translocation.^5,6^ More importantly, studies have increasingly supported the idea that presence or absence of EBV may represent a more clinically/molecularly relevant means of subtyping as opposed to traditional epidemiologic divisions.^7–10^

Known systematically as human herpesvirus 4, EBV has a double-stranded DNA genome measuring 172 kb.^11^ During initial infection of the epithelial cells, which primarily occurs via saliva,^12–15^ EBV has two primary stages: lytic and latent. In the lytic stage, the virus induces proliferation of the host epithelial cells in the salivary glands and produces infectious virions that can then infect other epithelial or B-cells. In the latent stage, the virus remains within the host B-cells with minimal viral protein production to avoid detection by the immune system but can gradually be replicated within host B-cells, leading to cell immortalization.^16–19^ The latent stage is most critical for tumorigenesis and has limited states of protein expression involving the latent membrane proteins (LMP1, LMP2A, LMP2B) and the EBV nuclear proteins (EBNA1, EBNA2, EBNA-3A, EBNA-3B, EBNA-3C, EBNA-LP), of which EBNA2 and EBNA3 show deep divergence classifying the virus into type 1 and type 2.^20–22^

Historically, studies have conformed to the dogma that EBV in BL is confined to the latency I state in which only the EBNA1 gene as well as BART and EBER miRNAs are expressed.^23,24^ Recent studies are finding, however, that heterogeneity across and within BL tumors may enable alternative latency states such as latency III where other genes including LMP1 are expressed.^25–29^ For example, Xue et al. identified LMP1 and LMP2A expression in BL tumors from Malawi.^29^ Willard and colleagues even reported spontaneous lytic BL cell lines that expressed LMP1.^25^

EBV type and variation are extensive and known to affect tumorigenesis.^30,31^ For example, whole genome sequencing (WGS) revealed from 101 BL tumors that viral presence drove mutations through the activation-induced cytidine deaminase (AID) mechanisms and defined a set of 72 driver genes, which were differentially expressed among the three epidemiologic subtypes of BL.^8^ Kaymaz et al. demonstrated that about a third of eBL tumors contain EBV type 2, whereas two-thirds are infected with EBV type 1 along with intertypic recombinants containing type 1 and type 2 regions.^9^ Kaymaz and colleagues further found that EBV type 1 tumors have fewer common mutations compared to type 2 and may attenuate the requirement for select driver mutations in BL tumorigenesis.^9^ Recent work has examined EBV in tumorigenesis and patient survival time, but so far it has focused on the major division of types 1 and 2. Thus, there is significant variation beyond type-specific variation that is currently unexplored. For example, Thomas et al. established genetic subtypes within BL based on the driver genes and compared EBV status and survival amongst their genetic subtypes but did not examine the relationship between driver genes and specific EBV variants.^32^ A recent study also found that some EBV-positive sBLs may be molecularly similar to EBV-positive eBLs.^33^ Thus, within EBV-positive BL, we sought to elucidate the role of specific EBV variants in affecting patient survival time, determine the association between specific EBV variants and driver genes in regards to patient survival time, and define new prognostic subtypes incorporating both EBV variants and driver genes.

## MATERIALS AND METHODS

### Study Population

Based on the known EBV statuses of the publicly available BL tumor WGS samples from Thomas et al. and Panea et al.,^8,32^ we downloaded only the EBV-positive samples as processed binary alignment map (BAM) files from the Burkitt Lymphoma Genome Sequencing Project (BLGSP)^10,32^ via the Genomic Data Commons (GDC) Data Portal (Project ID: CGCI-BLGSP, dbGaP Study Accession: phs000235) and our own publicly available samples from Panea et al.^8^ via the European Genome-Phenome Archive (Accession Number: EGAD00001005781) on August 31, 2023 (Figure 1). The BLGSP samples were obtained from patients in Uganda, the United States, France, Canada, and Brazil, while our samples were obtained from patients in Kenya (**Table 1**). Due to the growing shift away from traditional epidemiological subtypes, instead of categorizing patients as endemic or sporadic, patients were classified based on the country of origin’s income classification in the United Nations’ World Economic Situation and Prospects 2024.^34^ These categories included high-income countries including Canada, France, and the United States and low-middle-income countries including those classified as low-income (Uganda), lower-middle-income (Kenya), and upper-middle-income (Brazil).^35–37^

**Figure 1.**
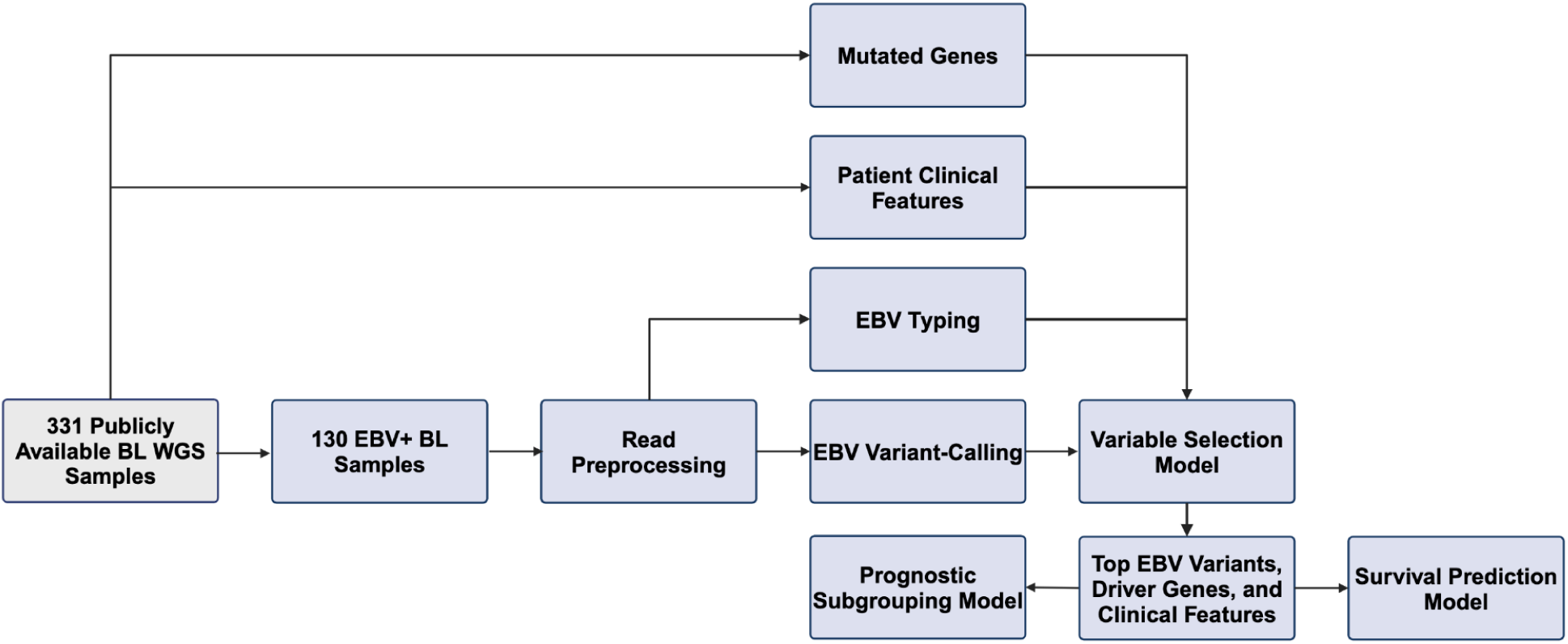
Analysis Pipeline. Out of the 331 original publicly available BL whole-genome-sequencing samples, sequencing reads from 130 EBV-positive samples were downloaded, preprocessed, typed for EBV, and called for EBV variation. Along with information on mutated genes and patient clinical features publicly available with the samples, EBV type and variation were inputted into a variable selection model to extract the top EBV variants, driver genes, and clinical features associated with patient survival time. These features were then applied into survival prediction and prognostic subgrouping models.

**Table 1.** Patient Characteristics.

|  | **No. Patients (%)** |
| --- | --- |
| **Sex** |  |
| Male | 87 (66.92) |
| Female | 43 (33.08) |
| **Median Age (range)** | 8.33 (0.94-90) |
| **EBV Type** |  |
| Type 1 | 103 (79.23) |
| Type 2 | 27 (20.77) |
| **Country Income Level** |  |
| High | 27 (20.77) |
| Low-Middle | 103 (79.23) |
| **Country** |  |
| Uganda | 70 (53.85) |
| Kenya | 25 (19.23) |
| United States | 16 (12.31) |
| Canada | 10 (7.69) |
| Brazil | 8 (6.15) |
| France | 1 (0.77) |

### Sequencing Read Preprocessing and Variant-Calling

To conduct a uniform process for variant-calling, all processed reads were trimmed of any residual adapters and converted to FASTQ. They were then mapped to the human (GRCh38), EBV1 (NC_007605), and EBV2 (NC_009334) reference genomes. EBV-specific reads were then preprocessed to remove read duplicates and sort uniformly with GATK 4.2.2.0 and SAMtools 1.16.1 per GATK best practices regarding data preprocessing for variant discovery (**Supplementary Figure 1**).^38,39,40^ Since the EBV strains from Thomas et al.’s samples were not typed, we uniformly typed all samples including those from Panea et al. with BEDtools and SAMtools based on the number of reads mapping to the *EBNA2* and *EBNA3* genes of the reference genomes for each type (**Supplementary Figure 2**).^40,41^

To determine the major EBV variants, the uniformly preprocessed reads across all samples were jointly variant-called using the EBV reference genome NC_007605 with three rounds of GATK4 HaplotypeCaller per GATK4’s best practices for germline short variant discovery akin to Kaymaz et al. albeit with haploid settings and revised functions based on the latest GATK updates (**Supplementary Figure 3**).^9,42^ SNPs with a read depth of less than 5 (DP<5), Phred-scaled probability that the site has no variant normalized by read depth of less than 2 (QD<2), mapping quality less than 10 (MQ<10), and strand bias using Fisher’s Exact Test greater than 60 (FS>60) were removed; while, indels with a read depth of less than 5 (DP<5), Phred-scaled probability that the site has no variant normalized by read depth of less than 2 (QD<2), and strand bias using Fisher’s Exact Test greater than 200 (FS>200) were removed. EBV variants were then annotated with SnpEff^43^ and filtered with SnpSift to remove synonymous EBV variants (**Supplementary Figure 4**).^44^ Each variant site was filtered to include only minor alleles >=5% and sites with <=2% missingness. EBV variants were further filtered to only include those in at least 5% of the 130 samples and sites missing in fewer than 2% of the samples (**Supplementary Figure 4**). Missing EBV variants for a given sample were imputed with the mode of the variant in question across all samples.

All publicly available samples had human somatic mutation data for recurrently mutated genes, hereon referred to as driver genes, previously defined and detected from their respective publications.^8,32^ Using this, somatic tumor mutation data in-common driver genes for the 130 WGS samples were extracted and preprocessed with in-house scripts. Then, for each sample, each driver gene was classified as mutated (1) if the sample had one or more missense, frameshift, nonsense, or splice site mutation or unmutated (0) otherwise. For each driver gene across all corresponding sites, driver genes were also filtered to only include genes for which were mutated (1) in at least 5% of the samples and data was missing in fewer than 2% (**Supplementary Figure 4**). Samples with missing data at a specific gene were imputed with the mode.

### Variable Selection and Survival Prediction Models

Along with clinical features including patient sex, age, EBV type, country, and country classification by income (high-income or low-middle-income), these filtered EBV variants and previously defined driver genes were then used as features in least absolute shrinkage and selection operator (LASSO) regularized and Bayesian variable selection models, both of which were implemented within a Cox proportional hazards framework with the glmnet^45^ and BVSNLP^46^ packages, respectively, in R^47^. (**Supplementary Figure 4**). These two variable selection approaches are commonly used to find the most important features in high-dimensional data where the number of samples n is significantly smaller than the number of features *p*. One advantage to implementing both models is that each tends to penalize correlated features differently. The LASSO will select a few representatives out of a group to be significant, while Bayesian shrinkage priors will penalize a group of features more evenly. By taking the consensus of both models, we are more likely to find a more robust set of EBV related features that are associated with survival.

Separate variable selection models were implemented for each of the following categories, hereinafter referred to as feature sets: 1) clinical features, 2) driver genes, 3) clinical features and driver genes, 4) EBV variants, 5) EBV variants and driver genes, 6) EBV variants and clinical features (EBV/clinical features model), and 7) EBV variants, driver genes, and clinical features (complete model). LASSO Cox and Bayesian Cox then determined which features were associated with patient survival time. For each feature set, features with both non-zero coefficients in LASSO Cox and posterior inclusion probabilities above zero in Bayesian Cox were considered the top features used for prediction and prognostic subgrouping. These top features were then used as the inputs to a Cox proportional hazards survival prediction model. Using the coxph package^48^ in R, each feature set’s prediction model was run over 1000 replicates with each iteration using a random 90-10 training-testing split of all samples. The variable selection and survival prediction models were implemented on both the overall cohort including all patients as well as the African cohort consisting of patients only from African countries (Kenya and Uganda). For each feature, the reported hazard ratio was the median value across all 1000 replicates. Predictive performances were measured with Harrell’s concordance index where a value of 1 corresponded to perfect prediction of which of any given two patients will die first across all possible pairs of patients, while a value of 0.5 was an uninformed prediction.^49^

### Prognostic Subgrouping Survival Model

Using the top features in the complete feature set, the optimal number of prognostic subgroups was determined with the silhouette score. The silhouette score measures the mean distance between points within a subgroup compared to the mean distance between points across neighboring subgroups. Final prognostic subgrouping was conducted with non-negative matrix factorization using the NMF package in R^50^, and Kaplan-Meier curves were performed to compare each prognostic subgroup’s overall survival.

### Data Availability

All coding scripts used in the pipelines (preprocessing, EBV typing, variant-calling, survival prediction, and prognostic subgrouping) are available on Github at https://github.com/bailey-lab/BL-Survival-Prediction-and-Prognostic-Subgrouping.

## RESULTS

### Identification of publicly available EBV-positive BL samples

Out of 331 publicly available BL tumor WGS samples from Thomas et al. (n=230) and Panea et al. (n=101),^8,32^ we extracted only the 105 and 25 EBV-positive samples, respectively, resulting in a final sample size of 130 EBV-positive BL samples (Figure 1). Of these 130 EBV-positive BL patients, 103 (79.2%) were from low-middle-income countries, while 27 (20.8%) were from high-income countries (**Table 1**). Countries of origin included Uganda (n=70, 53.8%), Kenya (n=25, 19.2%), the United States (n=16, 12.31%), Canada (n=10, 7.7%), Brazil (n=8, 6.15%), and France (n=1, 0.77%). The majority of patients were male (n=87, 66.9%), and 33.1% (n=43) were female with a median age of 8.3 years. EBV type 1 was the dominant type, representing 79.2% (n=103) of all genotyped samples, with type 2 being found in 14.8% and 22.3% of patients in high-income and low-middle-income countries, respectively. The median survival time for patients from low-middle-income countries was 1.9 years, while those from high-income countries did not have median survival times due to a greater than 50% survival rate.

### EBV coverage and variants

The median depth of coverage of the EBV genome across all 130 samples was 938.2. From an initial 10,347 EBV variants and 16,033 mutated genes, we extracted 724 EBV variants and 73 potential driver genes for further analysis after filtration (Methods).

### EBV/clinical features model on overall cohort found high number of LMP1 variants associated with patient survival time

Using an initial feature set of EBV variants and clinical features known as the EBV/clinical features model on the overall cohort, the variable selection models yielded 22 EBV variants including seven in LMP1 alone (Met129Ile/M129I, His101Gln/H101Q, Ile152Leu/I152L, Ile63Met/I63M, Gly212Ala/G212A, Gly212Ser/G212S, and Gly331Glu/G331Q) as most associated with patient survival time (**Table 2**). While only three out of the 22 EBV variants including Thr208Ala/T208A (BTRF1), Met12Leu/M12L (BDLF3), and Thr710Asn/T710N (BSLF1) were individually associated with patient survival time in the multivariable Cox proportional hazards analysis (**Table 2**), all 22 were used in the survival prediction model.

**Table 2.** Multivariable Cox Proportional Hazards Analysis of EBV Variants Associated with Overall Survival in EBV Feature Set.

| **EBV Variant (Gene)** | **Allele Frequency/N (%)** | **aHR (95% CI)** | **P** |
| --- | --- | --- | --- |
| Thr208Ala/T208A (BTRF1) | 123/130 (94.62) | 0.004 (0.000-0.061) | 8.095*10<sup>-5</sup> |
| Met12Leu/M12L (BDLF3) | 128/129 (99.22) | 0.017 (0.001-0.259) | 0.004 |
| Thr710Asn/T710N (BSLF1) | 16/130 (12.31) | 4.461 (1.133-17.574) | 0.033 |
| Met129Ile/M129I (LMP1) | 121/130 (93.08) | 35.888 (0.953-1318.905) | 0.053 |
| His101Gln/H101Q (LMP1) | 11/130 (8.46%) | 4.128 (0.945-17.792) | 0.060 |
| Ile152Leu/I152L (LMP1) | 55/129 (42.64) | 4.111 (0.738-23.153) | 0.106 |
| Ile63Met/I63M (LMP1) | 54/130 (41.54) | 0.143 (0.015-1.463) | 0.104 |
| Gly212Ala/G212A (LMP1) | 11/129 (8.53) | 0.380 (0.061-2.349) | 0.298 |
| Pro308Gln/P308Q (BVRF2) | 8/129 (6.20) | 3.120 (0.313-30.738) | 0.335 |
| Gly118Arg/G118R (BLLF1) | 15/130 (11.54) | 1.765 (0.557-5.559) | 0.337 |
| Val564_Ala568del/V564_A568del (EBNA-2) | 16/127 (12.60) | 0.454 (0.088-2.402) | 0.350 |
| Thr124Pro/T124P (BZLF1) | 39/129 (30.23) | 0.621 (0.200-1.925) | 0.407 |
| Pro622Ser/P622S (BBLF4) | 37/130 (28.46) | 1.348 (0.540-3.384) | 0.520 |
| Ala339Thr/A339T (BBLF4) | 122/129 (94.57) | 0.627 (0.086-4.583) | 0.643 |
| Arg648fs/R648fs (BHLF1) | 19/128 (14.84) | 1.236 (0.418-3.635) | 0.689 |
| Glu36Gln/E36Q (BNRF1) | 100/127 (78.74) | 0.792 (0.214-2.959) | 0.706 |
| Gly212Ser/G212S (LMP1) | 106/128 (82.81) | 1.218 (0.268-5.504) | 0.756 |
| Ala768Val/A768V (BLLF1) | 11/130 (8.46) | 1.130 (0.106-12.977) | 0.874 |
| Gly331Glu/G331Q (LMP1) | 7/129 (5.43) | 2.007*10<sup>-10</sup> (0.000-N/A) | 0.997 |
| Val479Ile/V479I (BRLF1) | 7/130 (5.38) | 7.003*10<sup>-8</sup> (0.000-N/A) | 0.998 |
| Val611Ile/V611I (BBLF4) | 37/130 (28.46) | N/A | N/A |
| Pro486Ser/P486S (BRLF1) | 7/130 (5.38) | N/A | N/A |
| **Country Income Level** |  |  |  |
| Low-Middle | 103 (79.23) | 12.099 (1.318-104.407) | 0.027 |

### EBV/clinical features model on African cohort still indicated a high number of significant variants in LMP1 but not EBV type

When limited to patients in Africa (n=95) for the African cohort, six of the 22 EBV variants from the overall cohort remained significantly associated with patient survival time (P308Q, I63M, H101Q, A339T, I152L, and G212A) with four of those coming from LMP1 alone (I63M, H101Q, I152L, and G212A) (**Supplementary Table 1**). In this variable selection model, six of the 19 variants significantly associated with patient survival time were located in LMP1 (Leu25Ile/L25I, I63M, H101Q, I152L, G212A, Gln322His/Q322H). EBV type was still not significantly associated with patient survival time, and while there were three significant variants in EBNA2 in the initial analysis, removing all type-specific variants did not alter the lack of association between EBV type and patient survival time.

### Country income level was the strongest individual feature among all features associated with patient survival time but EBV type was not associated

In the EBV/clinical features model on the overall cohort, country income level was the single most significant feature of all features associated with patient survival time with a hazard ratio of 12.10 (**Table 2**). Country level income was also the only clinical feature associated with patient survival time, as EBV type, patient sex, age, and country were not significantly associated. When limited to African patients only, there was no statistically significant survival difference between patients in Kenya (n=25) and patients in Uganda (n=70, p=0.44). Removing type-specific variants such as those within the EBNA2 and EBNA3 genes did not change the lack of association.

### Complete model on overall cohort found eight tumor driver genes and nine EBV variants including four in LMP1 associated with patient survival time

In the complete model, which consisted of all clinical features, tumor driver genes, and EBV variants, the variable selection models identified eight driver genes including CNTNAP3B, ID3, HIST1H2BK, MAP3K9, IGK, ETS1, P2RY8, and TP53 in addition to nine EBV variants with four in LMP1 as most associated with patient survival time (**Table 3**). While only two driver genes had independent statistically significant hazard ratios (CNTNAP3B: adjusted hazard ratio 4.03, 95% CI 1.17-14.03, p=0.03; ID3: aHR 0.42, 95% CI 0.19-0.95, p=0.04), all eight were used in the survival prediction model. This model also uncovered nine EBV variants including four LMP1 variants (H101Q, I152L, G212A, and G331Q) as most associated with patient survival time, indicating that 13 of the 22 variants from the EBV variant-only feature set were no longer significant after incorporating driver genes. Cox proportional hazards analysis identified six of the nine EBV variants as individually associated with patient survival time (M12L/BDLF3: aHR 2.00*10^-3^, 95% CI 0.00-0.04, p=2.63*10^-9^; H101Q/LMP1: aHR 10.94, 95% CI 3.27-36.86, p=9.93*10^-5^; T208A/BTRF1: aHR 4.00*10^-3^, 95% CI 0.00-0.07, p=2.42*10^-4^; T710N/BSLF1: aHR 3.89, 95% CI 1.61-9.40, p<0.01; Pro308Gln/P308Q/BVRF2: aHR 4.17, 95% CI 1.23-13.35, p=0.02; I152L/LMP1: aHR 2.43, 95% CI 1.18-5.01, p=0.02). A complete feature set analysis on the African cohort shared three of the nine EBV variants (P308Q, H101Q, and I152L) and three of the eight driver genes (HIST1H2BK, ETS1, and ID3) in common with the overall cohort (**Supplementary Table 2**).

**Table 3.** Multivariable Cox Proportional Hazards Analysis of driver genes, EBV Variants, and Clinical Features Associated with Overall Survival in Complete Feature Set.

| EBV Variant (Gene) | Allele Frequency/N (%) | aHR (95% CI) | P |
| --- | --- | --- | --- |
| Met12Leu/M12L (BDLF3) | 128/129 (99.22) | 0.002 (0.000-0.036) | 2.625*10<sup>-9</sup> |
| His101Gln/H101Q (LMP1) | 11/130 (8.46%) | 10.942 (3.273-36.860) | 9.929*10<sup>-5</sup> |
| Thr208Ala/T208A (BTRF1) | 123/130 (94.62) | 0.004 (0.000-0.074) | 2.418*10<sup>-4</sup> |
| Thr710Asn/T710N (BSLF1) | 16/130 (12.31) | 3.887 (1.614-9.395) | 0.002 |
| Pro308Gln/P308Q (BVRF2) | 8/129 (6.20) | 4.173 (1.230-13.348) | 0.016 |
| Ile152Leu/I152L (LMP1) | 55/129 (42.64) | 2.434 (1.181-5.009) | 0.016 |
| Gly212Ala/G212A (LMP1) | 11/129 (8.53) | 0.260 (0.033-2.182) | 0.222 |
| Val564_Ala568del/V564_A568del (EBNA-2) | 16/127 (12.60) | 0.653 (0.136-3.234) | 0.602 |
| Gly331Glu/G331Q (LMP1) | 7/129 (5.43) | 1.515*10<sup>-9</sup> (0-N/A) | 0.997 |
| **Driver Gene** |  |  |  |
| CNTNAP3B | 16/130 (12.31) | 4.025 (1.172-14.025) | 0.027 |
| ID3 | 55/130 (42.31) | 0.424 (0.189-0.946) | 0.036 |
| HIST1H2BK | 14/130 (10.77) | 2.329 (0.851-6.398) | 0.100 |
| MAP3K9 | 12/130 (9.23) | 0.194 (0.017-2.372) | 0.204 |
| IGK | 33/130 (25.38) | 0.559 (0.222-1.405) | 0.217 |
| ETS1 | 17/130 (13.08) | 1.559 (0.608-4.035) | 0.352 |
| P2RY8 | 10/130 (7.69) | 0.928 (0.097-3.387) | 0.520 |
| TP53 | 25/130 (19.23) | 0.926 (0.330-2.610) | 0.797 |
| **Country Income Level** |  |  |  |
| Low-Middle | 103 (79.23) | 37.166 (3.548-373.040) | 0.002 |

### Interactions between driver genes, specific EBV variants, and country income level required for best survival predictive performance

To better understand variable selection across features, we explored additional models, examining a total of seven variable selection models (clinical features, driver genes, clinical features/driver genes, EBV variants, EBV variants/driver genes, EBV variants/clinical features, and EBV variants/driver genes/clinical features). For each, we incorporated the corresponding features to predict survival and compared their performance. Each model had the following number of features: clinical features=1, driver genes=33, clinical features/driver genes=34, EBV variants=22, EBV variants/driver genes=16, EBV variants/clinical features=23, and EBV variants/driver genes/clinical features=18. Since EBV type, patient sex, age, and country were not associated with survival from the variable selection models, these clinical features were not applied in the survival prediction models. The survival model with the lowest predictive performance used country income level as its only feature and had a testing concordance index (c-index) of 0.56 (**Table 4**). The next best performing variable selection and survival prediction model only used driver genes in its feature set and had a testing c-index of 0.57 (p=0.11 compared to country income level-only model). When both driver genes and country income level were used for the feature set, the model had an improved testing c-index of 0.61 (p<0.01 compared to driver gene-only model). Compared to the model with driver genes and country income level, the model trained with EBV variants only had an even better testing c-index of 0.72 (p<0.01 compared to driver gene + country income level model). The model using both EBV variants and driver genes performed similarly with a testing c-index of 0.72 (p=0.39). In comparison, the model with EBV variants and country income level had an improved testing c-index of 0.73 (p=0.04). Finally, a model trained on the complete feature set of EBV variants, driver genes, and country income level had the best testing c-index at 0.76 (p<0.01), which was significantly higher than all prior models, indicating that interactions between EBV variants, driver genes, and clinical features were required for the best survival predictive performance.

**Table 4.** Survival Prediction Model Performances.

| Feature Set | Training Set C-Index | Testing Set C-Index | p-value* |
| --- | --- | --- | --- |
| Complete (EBV + Tumor + Clinical Features) | 0.8504 ± 0.0003 | 0.7572 ± 0.0038 | < 0.0001 |
| EBV + Clinical Features | 0.8356 ± 0.0003 | 0.7324 ± 0.0040 | 0.0358 |
| EBV + Tumor | 0.7968 ± 0.0004 | 0.7226 ± 0.0038 | 0.3912 |
| EBV | 0.8184 ± 0.0005 | 0.7210 ± 0.0041 | < 0.0001 |
| Tumor + Clinical Features | 0.8144 ± 0.0005 | 0.6101 ± 0.0047 | < 0.0001 |
| Tumor | 0.7881 ± 0.0005 | 0.5739 ± 0.0046 | 0.1098 |
| Clinical Features | 0.6085 ± 0.0005 | 0.5601 ± 0.0124 | N/A |

\*T-test comparing testing set C-index of current feature set relative to that of next-best performing feature set

When limited to African patients only, the EBV/clinical features model had a testing c-index of 0.67. The complete model on the African cohort observed a performance boost to 0.71 for the testing c-index.

### Three prognostic subgroups were defined with differential survival

In order to explore patterns and potential groupings of features relative to survival, we employed silhouette analysis to determine the optimal number of subgroups and non-negative matrix factorization to cluster features into meta-features and define prognostic subgroups that may have differential survival rates. Using the complete feature set of EBV variants, driver genes, and clinical features, the silhouette score indicated that three prognostic subgroups yielded the highest cluster stability. These subgroups were defined by different meta-features, which were composed of various driver genes and EBV variants (Figure 2). Subgroup 1 was defined by a meta-feature composed of driver genes TP53, CNTNAP3B, MAP3K9, P2RY8, and IGK and the EBV variants G331Q (LMP1) and H101Q (LMP1) (Figure 3). Subgroup 2 was defined by a meta-feature consisting of the driver gene ID3 and a different LMP1 variant I152L (LMP1) and EBV variant T710N (BSLF1). Finally, Subgroup 3 was defined by a different meta-feature that included the driver genes ETS1 and HIST1H2BK as well as a different LMP1 variant G212A and EBV variants Val564_Ala568del/V564_A568del (EBNA-2), P308Q (BVRF2), M12L (BDLF3), and T208A (BTRF1). In a Kaplan-Meier survival analysis, Subgroup 1 observed the highest overall survival, followed by Subgroup 2, and then finally Subgroup 3 (p=0.03, Figure 4).

**Figure 2.**
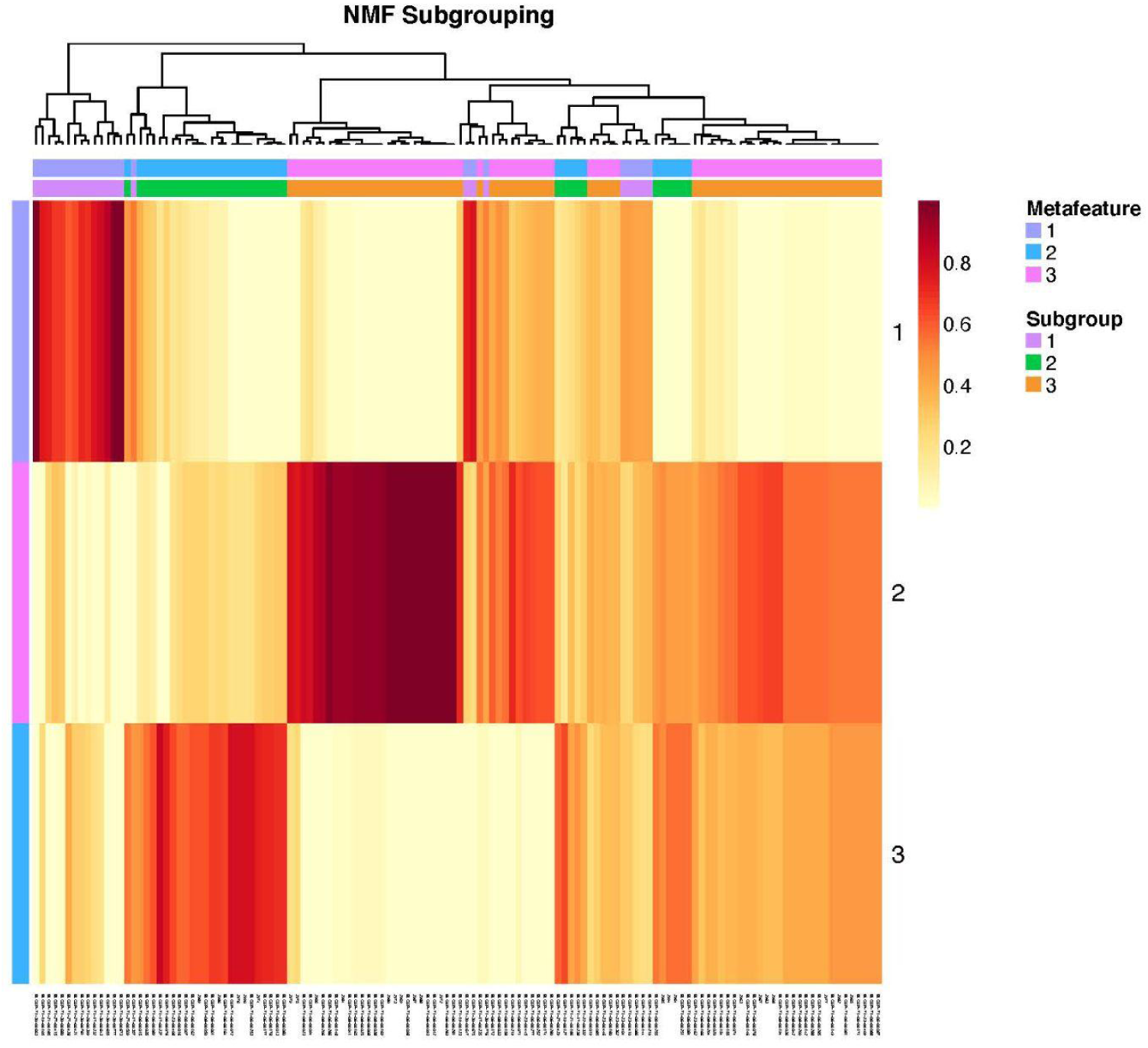
Prognostic Subgroups with Dominant Metafeatures. Based on the silhouette score, all samples were categorized into one of three prognostic subgroups. The columns represent the samples, and the two color schemes correspond to the metafeature number and prognostic subgroup number. All samples and their subgroups are shown here in this hierarchical clustering.

**Figure 3.**
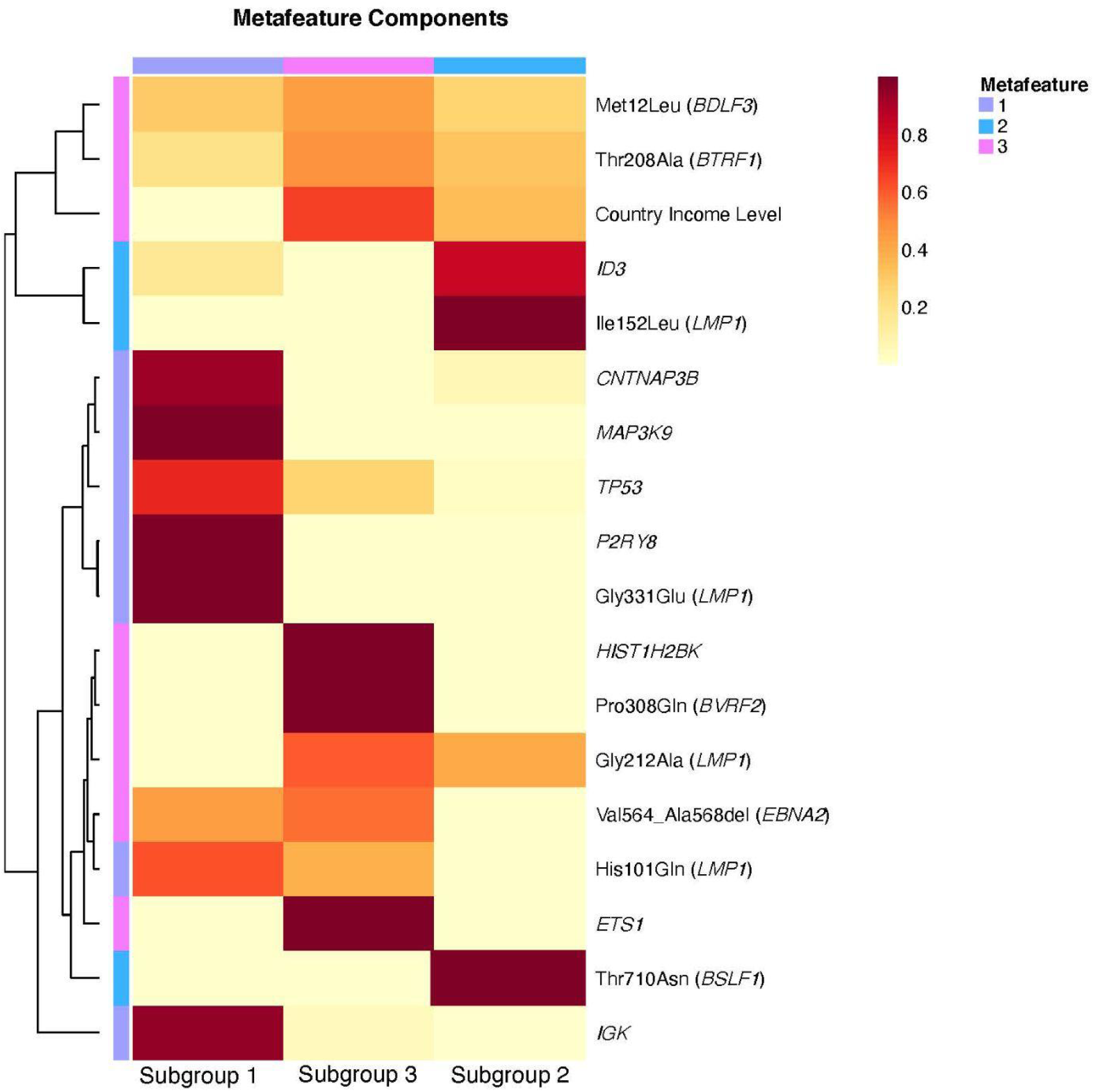
EBV Variants/Driver Genes/Country Income Level Contributions to Metafeatures. Each metafeature was defined by different dominant genes. Metafeature 1 consisted of CNTNAP3B, MAP3K9, TP53, P2RY8, IGK, G331Q (LMP1), and H101Q (LMP1). Metafeature 2 was defined by ETS1, HIST1H2BK, G212A (LMP1), V564_A568del (EBNA2), P308Q (BVRF2), M12L (BDLF3), and T208A (BTRF1). Finally, Metafeature 3 was composed of ID3, I152L (LMP1), and T710N (BSLF1). Note that all three metafeatures included a different LMP1 variant.

**Figure 4.**
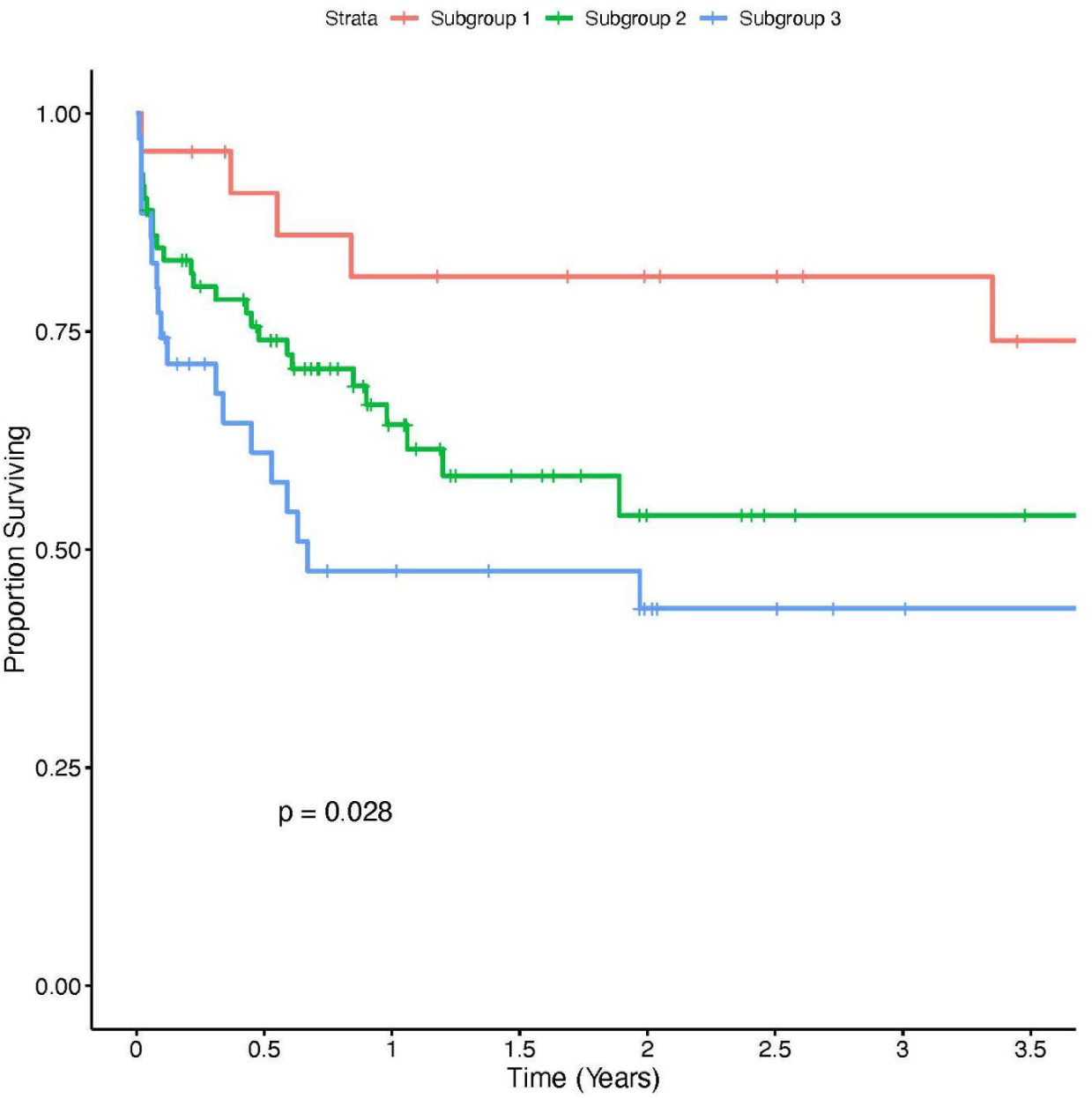
Overall Survival of Prognostic Subgroups. The prognostic subgroups observed differential survival with patients from Subgroup 1 experiencing the best survival, followed by those in Subgroup 2, and finally the patients in Subgroup 3.

These prognostic subgroups included geographical differences (p<0.01). For example, Subgroup 1 was predominantly composed of North American and European patients (78.26%, n=18), while Subgroups 2 and 3 had higher proportions of African and South American patients (Subgroup 2: 97.22%, n=70 and Subgroup 3: 80.00%, n=28). Notably, geography was not absolute amongst these subgroups, meaning that there were some African and South American patients in Subgroup 1 (21.74%, n=5) and North American and European patients (Subgroup 2: 2.78%, n=2 and Subgroup 3: 20.00%, n=7) in Subgroups 2 and 3 (**Supplementary Table 3**).

## DISCUSSION

BL is an aggressive B-cell lymphoma that is fatal if left untreated. While the prognosis of BL is known to depend on factors such as the stage of the disease, other molecular and genetic interactions including viral variation have not been comprehensively explored. In this study, we incorporated EBV variation along with clinical features and driver gene mutations into survival models for patients with EBV-positive tumors. EBV variants, particularly those in LMP1, were found to be associated with survival. Not surprisingly, we found that study participants’ country income level (a likely surrogate for clinical access and care) was the strongest individual feature associated with BL patient survival time. An analysis limited to patients in Africa shared many of the same variants as that of the complete cohort and still found a disproportionately higher number of significant variants in LMP1, but not EBV type to be associated with patient survival time, indicating that the addition of patients outside of Africa did not drastically alter our models. We also defined potential interactions between specific EBV variants and driver genes in the context of patient survival time. Using the associated EBV variants, clinical features, driver genes, we defined three new prognostic subgroups that showed differing survival.

The EBV variants associated with patient survival were especially enriched for LMP1 variants. LMP1 is the the main transforming oncoprotein in EBV and has a range of biological roles including inhibiting apoptosis and promoting cell proliferation through activation of the NF-kB pathway.^51^ Amino acid 212 in LMP1 was especially intriguing, as G212S had a hazard ratio greater than 1, while G212A had a hazard ratio less than 1. A prior study demonstrated that the G212S mutation in EBV-positive post-transplant lymphoproliferative disorder (PTLD) B-cell lines was associated with increased extracellular signal-regulated kinase (ERK)/mitogen-activated protein kinases (MAPK) activation and c-Fos expression, indicating this variant may be a gain-of-function mutation.^52^ Furthermore, a clinical trial of 872 pediatric transplant recipients by Martinez et al. reported that G212S increased the risk of EBV-positive PTLD by nearly 12-fold.^53^ While no studies have investigated the functional role of a substitution to alanine in place of serine, Martinez et al. found that 0% of EBV-positive PTLD patients had G212A but 96.9% carried G212S,^53^ suggesting that this non-synonymous substitution may significantly dampen the virus’s ability for tumorigenesis and aggressiveness of the resulting tumor. Despite the traditional dogma that BL is classically latency I, our findings support those of more recent studies highlighting the expression and potential significance of LMP1 in BL ^25,29^ and highlight the potential for using specific EBV variants such as G212S and G212A as predictive biomarkers in EBV-positive BL (**Figure 5**).

**Figure 5.**
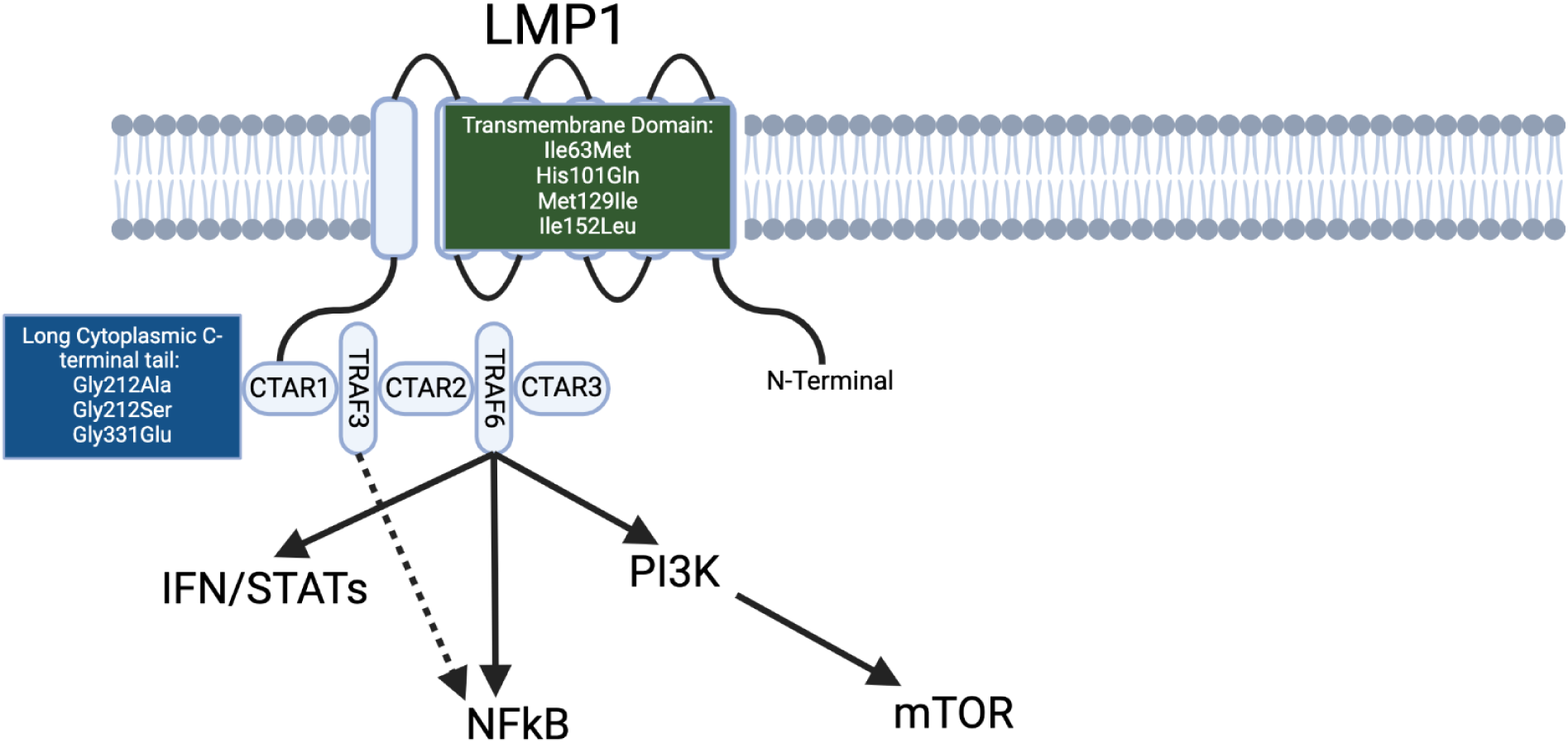
Model of LMP1 Pathways and Variants. Four out of the seven significant LMP1 variants (Ile633Met, His101Gln, Met129Ile, and Ile152Leu) corresponded to the transmembrane domain, while the other three (Gly212Ala, Gly212Ser, and Gly331Glu) belonged to the domain of the long cytoplasmic C-terminal tail, which is involved in the NFkB and PI3K/mTOR pathways. Adapted from sources.^69–71^

When we restricted our variable selection models to patients in Africa, we identified 6 of the same 22 EBV variants from the original cohort and still found a lack of association between EBV type and survival time, indicating that the incorporation of patients outside of Africa did not completely change our model and supporting the growing number of studies that propose traditional epidemiologic subtypes corresponding to patient geography may not be the best classification of BL.^7–9^ Importantly, we continued to find a high number of variants associated with patient survival time in the LMP1 gene including G212A and I152L and a lack of association between EBV type and patient survival time despite prior studies showing molecular differences between Types 1 and 2 in the context of tumorigenesis. For example, Panea et al. and Agwati et al. reported that EBV type 1 tumors have a higher mutational load compared to EBV type 2.^8,54^ It is also known that Type 1 more effectively immortalizes B-cells^55^ and is more prevalent in endemic BL patients compared to healthy controls in Africa.^56^ Based on our findings, however, it seems that these differences do not translate to differential patient survival time.

Of all features, country income level was the most significant contributor to patient survival time with patients in low-middle-countries experiencing the highest hazard ratios in all relevant feature sets. In this study, patients from low-middle-income countries, namely countries on the continents of Africa and South America, had a median survival time of 1.89 years compared to those from high-income countries which observed a greater than 50% survival rate. This finding supports those of many prior studies, as BL patients in equatorial Africa have a survival rate nearly half of that of BL in non-tropical countries.^4^ Country income level is essentially a proxy for access to healthcare, and since BL is a highly aggressive tumor that requires intensive and consistent treatment, limited access to care such as delays in diagnosis and insufficient clinical supportive care can have potentially devastating consequences. For example, BL patients in sub-Saharan Africa are often diagnosed simply based on clinical presentation or tumor morphology without proper staging via microscopy of cerebrospinal fluid.^57^ These diagnostic methods can potentially miss some patients, leading to delays in diagnosis and advanced disease presentation. Altogether, our findings suggest that while genetic features of the human tumor cells and virus potentially underlying patient geography play a significant role in patient survival time, socioeconomic factors are by far the most important contributor.

Besides EBV variation, our variable selection and survival prediction models highlighted eight driver genes, two of which were individually associated with patient survival time as supported by prior studies.^8,32,58,59^ Contactin-associated protein family member 3B (CNTNAP3B), which was associated with decreased patient survival time, plays an integral role in cell adhesion, and abnormalities in the gene have been implicated in other cancers such as B-cell acute lymphoblastic leukemia.^58^ Furthermore, due to its association with immune checkpoint expression, CNTNAP3B has been used in a prognostic model for bladder cancer.^59^ ID3 was associated with increased patient survival time, and Thomas et al. similarly reported that patients with ID3 mutations observed increased survival compared to those who did not when controlling for TP53 mutations.^32^ Panea et al. found that ID3 was one of the most commonly silenced genes, and in vivo knockout of ID3 amplified the tumorigenic effects of MYC.^8^ Thus, while ID3 mutations may play a key role in the initial tumorigenesis, it could lead to less aggressive disease once the tumor has been established. Our results do differ with those of some prior studies. For instance, previous studies found that BL patients with TP53 mutations were more likely to relapse or experienced worse survival.^60,61^ While our multivariable Cox proportional hazards analysis did not find TP53 mutations to have a statistically significant association with patient survival time, our cohort consisted of EBV-positive BL patients, most of whom originated from Africa and South America, while most prior studies studied both EBV-negative and EBV-positive patients mostly in North America or Europe. Thus, given that EBV infection alters the driver genes in BL tumorigenesis, TP53 may not be significantly associated with clinical outcome in EBV-positive BL patients. Regardless, interactions between all three feature sets including driver genes, various specific EBV variants, and country income level were required for the best possible survival predictive performance.

Having defined interactions between specific EBV variants and driver genes, we found clinically relevant prognostic subgroups that demonstrated differences in survival and included a different LMP1 variant and various driver genes in each subgroup. Subgroup 1 consisted of the LMP1 variants Gly331Glu and His101Gln along with the driver genes TP53 and MAP3K9. Wang and colleagues determined that the TP53-encoded p53 protein activates LMP1, which in turn blocks many effects of p53 during viral transformation and enables EBV-infected cells to endure cellular damages.^62^ LMP1 is also known to regulate MAPK signal pathways.^52,63^ Subgroup 2 included the LMP1 variant I152L and driver gene ID3. ID3 is a well-characterized tumor suppressor gene in BL that amplifies the effects of MYC, thereby causing tumorigenesis.^8^ Recent studies have reported increased methylation and silencing of ID3 in EBV-infected B-cells via an LMP1-involved mechanism.^64^ Lastly, Subgroup 3 was composed of the LMP1 variant G212A, EBNA2 variant V564_A568del, and the driver gene ETS1. Kim et al. reported that LMP1 induced the expression of ETS1, thereby increasing the invasiveness of the tumor in nasopharyngeal carcinoma.^65^ Zhang and colleagues found that LMP1 and EBNA2, which is critically involved in latent viral transcription, comprise the minimum set of EBV genes required for tumorigenesis.^66,67^ On a larger cohort of both EBV-negative and EBV-positive BL samples, Thomas et al. also defined three subgroups using tumor mutations only. Given that most of our samples came from Thomas et al, their subgroups shared significant similarities yet had some key differences. For example, two of their three subgroups were defined by the same key driver genes: ID3 and TP53.^32^ However, unlike our subgroups, their subgroups did not incorporate viral variation and did not observe statistically significant differences in survival. With recent studies indicating that EBV status may be more defining than epidemiologic subtype,^8^ our findings suggest the presence of subgroups within EBV-positive BL with each having a different prognosis. Altogether, our findings portend the significance of the virus and its variants in impacting patient survival time and prognostic subgrouping.

Overall, our study identified specific EBV variants significantly associated with BL patient survival time. The main limitation of this study was its small sample size, as only 130 out of the 331 publicly available BL samples were EBV-positive, of which 103 were African or South American. However, our variable selection models were designed to address this limitation by decreasing the number of possible features associated with patient survival. As a result, we found a high number of significant variants in the LMP1 gene and lack of association with EBV type, findings which were confirmed in a sub-analysis of only patients from Africa. We determined that country income level, which corresponds to access to healthcare, was the most important factor in patient survival time. We also described potential interactions between driver genes and specific EBV variants in BL in the context of patient survival time and proposed a model that accurately predicts survival based on clinical features, driver genes, and EBV variants as well as defined prognostic subgroups. While prior studies have investigated the relationship between patient survival time and clinical features as well as driver genes,^4,68^ none have examined the role of EBV variation and its interactions with the aforementioned factors.

## CONCLUSIONS

Our study offers key insight into the role of specific EBV variants in BL patient survival time and their interactions with driver genes and clinical features, potentially introducing specific EBV variants as predictive biomarker candidates in future studies. We also define a survival prediction model and prognostic subgroups that may be used to assess prognosis for EBV-positive BL patients. Validation of these findings on larger patient cohorts as they become available are warranted.

## Acknowledgements

We thank Dr. Abebe Fola for his thoughtful comments.

