## Supplemental Figures 1-4, Supplemental Tables 1-3, Supplemental Methods for "Incorporation of Epstein-Barr viral variation implicates significance of LMP1 in survival prediction and prognostic subgrouping in Burkitt lymphoma"

**Supplementary Figure 1**. Read Preprocessing Pipeline

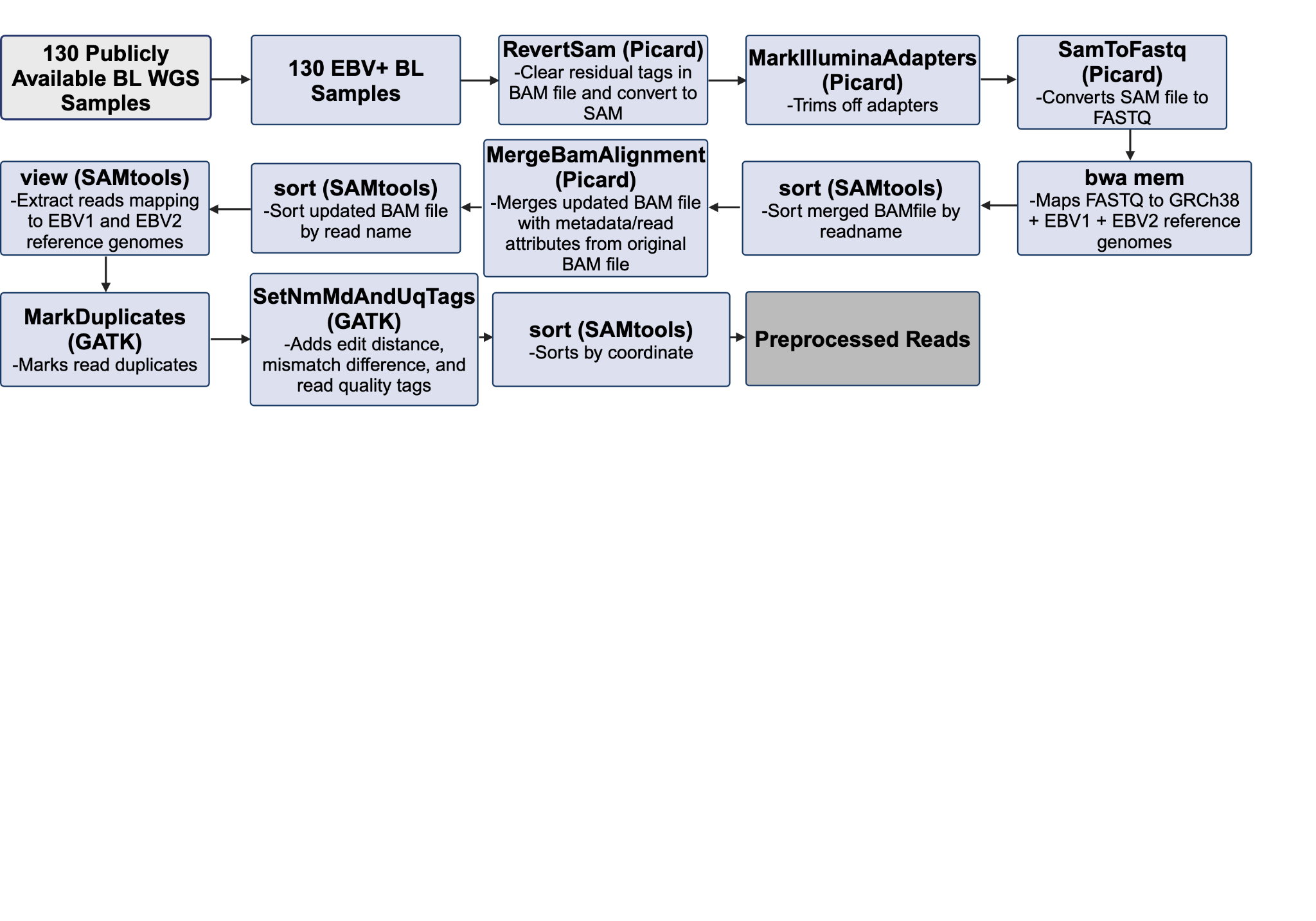

**Supplementary Figure 2**. EBV Typing Pipeline

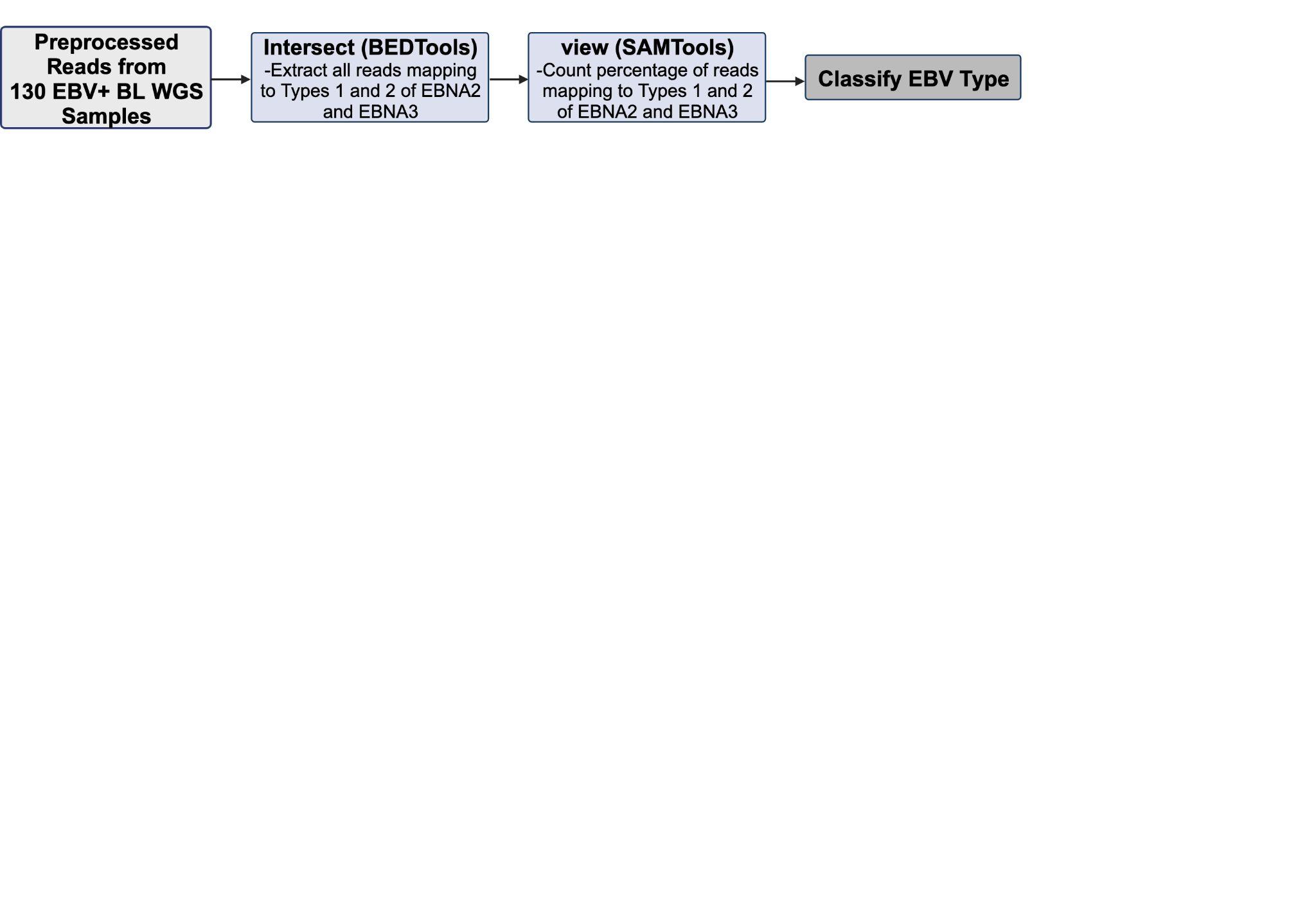

**Supplementary Figure 3**. EBV Variant-Calling Pipeline

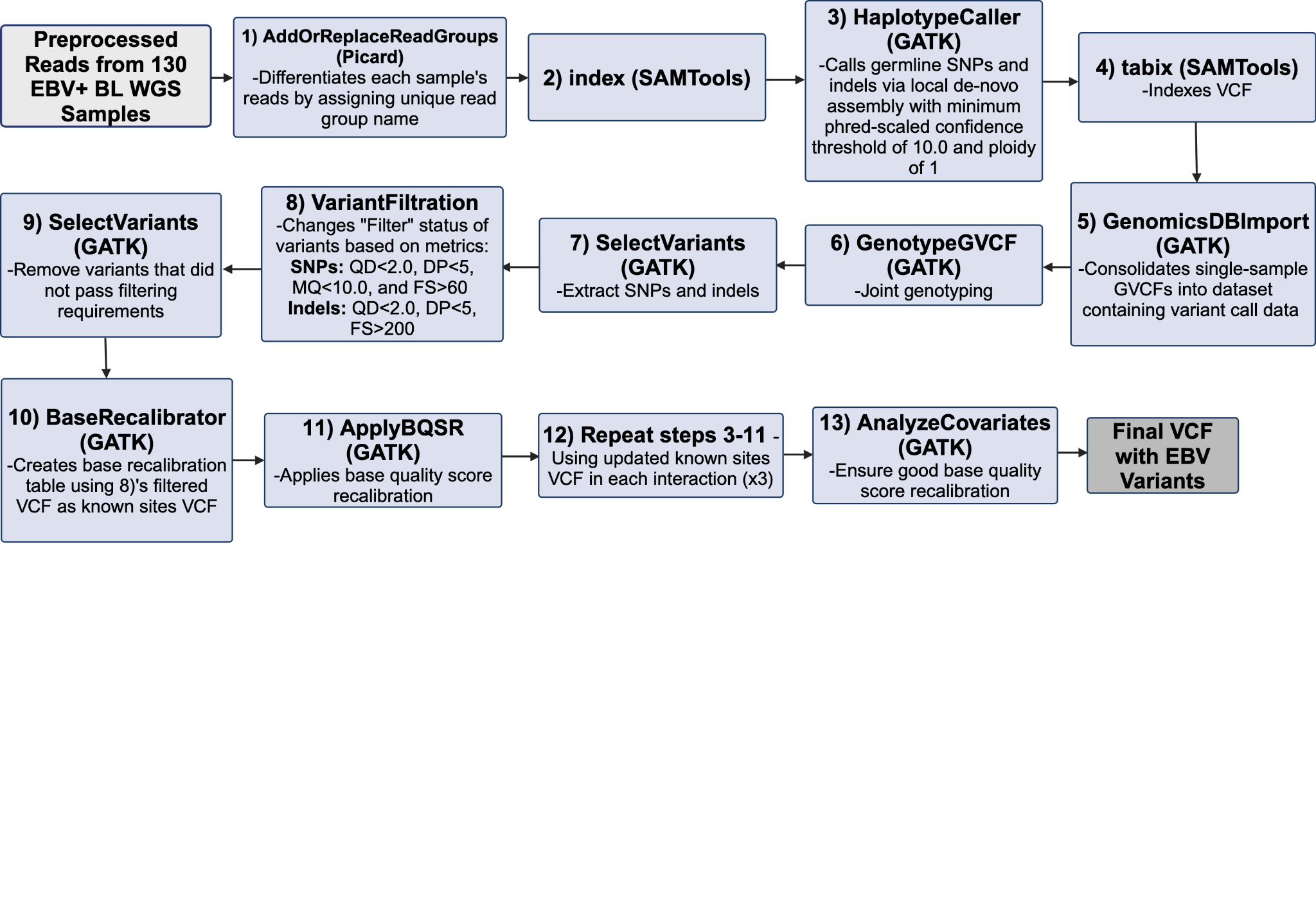

**Supplementary Figure 4**. Survival Prediction and Prognostic Subgrouping Pipeline

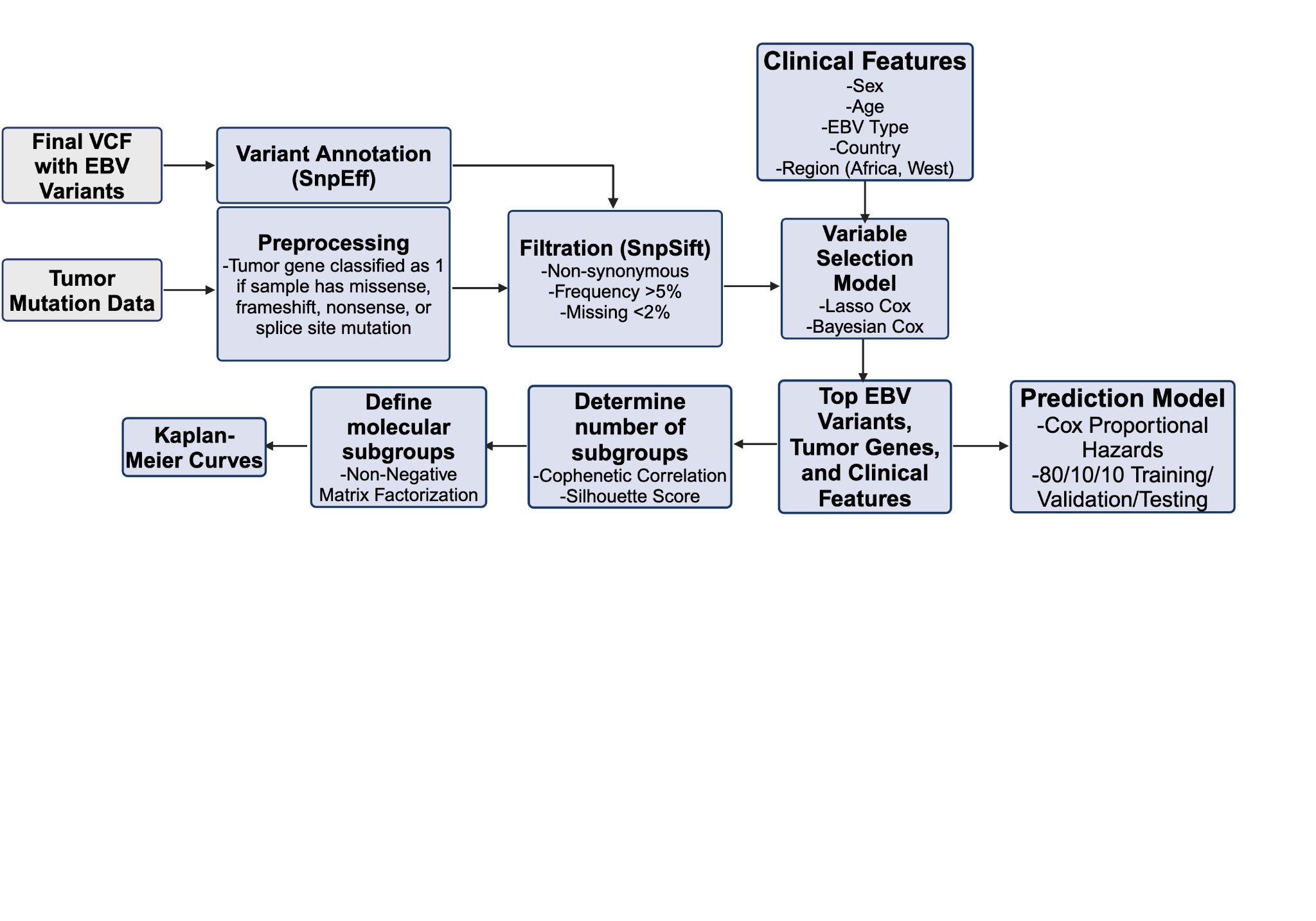

### SUPPLEMENTAL TABLES

**Supplementary Table 1****. Multivariable Cox Proportional Hazards Analysis of EBV Variants Associated with Overall Survival in EBV Feature Set for African Patients Only**

| **EBV Variant (Gene)** | **Allele Frequency/N (%)** | **aHR (95% CI)** | ***P*** |
| --- | --- | --- | --- |
| Pro308Gln/P308Q (BVRF2) | 8/94 (8.51) | 5.283 (1.721-16.349) | 0.004 |
| Ile115Thr/I115T (EBNA2) | 17/93 (18.28) | 4.813 (0.828-25.978) | 0.082 |
| Leu25Ile/L25I (LMP1) | 42/94 (44.68) | 0.115 (0.008-1.659) | 0.113 |
| Asp287Glu/D287E (EBNA2) | 18/94 (19.15) | 0.293 (0.072-1.312) | 0.113 |
| Ile63Met/I63M (LMP1) | 39/95 (41.05) | 7.472 (0.621-88.806) | 0.114 |
| His101Gln/H101Q (LMP1) | 8/95 (8.42) | 3.153 (0.716-14.201) | 0.131 |
| Asn302Ser/N302S (BNRF1) | 87/93 (93.55) | 0.388 (0.098-1.507) | 0.170 |
| Pro514dup/P514dup (LMP2A) | 25/95 (26.32) | 0.528 (0.206-1.354 | 0.186 |
| Lys586Arg/K586R (EBNA1) | 17/95 (17.89) | 0.470 (0.136-1.643) | 0.238 |
| Ala339Thr/A339T (BBLF4) | 89/94 (94.68) | 0.297 (0.035-1.435) | 0.258 |
| Ile152Leu/I152L (LMP1) | 41/94 (43.62) | 3.031 (0.321-29.146) | 0.324 |
| Gly212Ala/G212A (LMP1) | 10/95 (10.53) | 0.244 (0.020-3.545) | 0.324 |
| Cys23fs/C23fs (BHLF1) | 11/93 (11.83) | 1.885 (0.489-7.321) | 0.351 |
| Gln214Glu/Q214E (BARF0) | 27/94 (28.72) | 1.689 (0.552-5.140) | 0.360 |
| Ala151Thr/A151T (BOLF1) | 23/95 (24.21) | 0.917 (0.289-2.963) | 0.716 |
| Gln322His/Q322H (LMP1) | 33/95 (34.74) | 0.974 (0.123-7.811) | 0.719 |
| Ser10Gly/S10G (BOLF1) | 43/95 (45.26) | 1.087 (0.329-3.581) | 0.774 |
| Pro513fs/P513fs (LMP2A) | 6/95 (6.32) | 3.173*10^-8^ (0.000-N/A) | 0.997 |
| Arg114His/R114H (EBNA2) | 17/93 (18.28) | N/A | N/A |

**Supplementary Table 2****. Multivariable Cox Proportional Hazards Analysis of driver genes, EBV Variants, and Clinical Features Associated with Overall Survival in Complete Feature Set for African Patients Only**

| **EBV Variant (Gene)** | **Allele Frequency/N (%)** | **aHR (95% CI)** | ***P*** |
| --- | --- | --- | --- |
| Pro308Gln/P308Q (BVRF2) | 8/94 (8.51) | 10.521 (2.738-41.038) | 0.001 |
| His101Gln/H101Q (LMP1) | 8/95 (8.42) | 13.564 (1.833-103.814) | 0.011 |
| Pro514dup/P514dup (LMP2A) | 25/95 (26.32) | 0.273 (0.087-0.840) | 0.023 |
| Arg114His/R114H (EBNA2) | 17/93 (18.28) | 2.435 (0.762-7.817) | 0.135 |
| Lys586Arg/K586R (EBNA1) | 17/95 (17.89) | 0.370 (0.082-1.699) | 0.196 |
| Cys23fs/C23fs (BHLF1) | 11/93 (11.83) | 2.161 (0.569-8.306) | 0.256 |
| Ile152Leu/I152L (LMP1) | 41/94 (43.62) | 2.782 (0.474-16.258) | 0.259 |
| Gln322His/Q322H (LMP1) | 33/95 (34.74) | 2.150 (0.393-11.720) | 0.376 |
| Leu25Ile/L25I (LMP1) | 42/94 (44.68) | 0.534 (0.085-3.322) | 0.498 |
| Asn302Ser/N302S (BNRF1) | 87/93 (93.55) | 0.691 (0.158-3.059) | 0.585 |
| Ala339Thr/A339T (BBLF4) | 89/94 (94.68) | 0.927 (0.089-9.529) | 0.758 |
| Pro513fs/P513fs (LMP2A) | 6/95 (6.32) | 3.947*10^-8^ (0.000-N/A) | 0.997 |
| Ile115Thr/I115T (EBNA2) | 17/93 (18.28) | N/A | N/A |
| **Driver Gene** |  |  |  |
| IGL | 63/95 (66.32) | 0.332 (0.133-0.821) | 0.017 |
| PCLO | 5/95 (5.26) | 5.464 (0.945-32.491) | 0.058 |
| HIST1H2BK | 12/95 (12.63) | 2.857 (0.914-8.885) | 0.071 |
| GNAI2 | 12/95 (12.63) | 2.388 (0.809-7.079) | 0.114 |
| ETS1 | 16/95 (16.84) | 1.805 (0.644-5.075) | 0.261 |
| ID3 | 42/95 (44.21) | 0.592 (0.232-1.515) | 0.276 |
| SIN3A | 16/95 (16.84) | 1.217 (0.339-4.429) | 0.678 |
| HIST1H1C | 7/95 (7.37) | 1.243 (0.307-5.176) | 0.720 |

**Supplementary Table 3**. Geographical Distribution Among Prognostic Subgroups

| **Subgroup** | **No. North American/European Patients (%)** | **No. African/South American Patients (%)** |
| --- | --- | --- |
| 1 | 18 (78.26) | 5 (21.74) |
| 2 | 2 (2.78) | 70 (97.22) |
| 3 | 7 (20.00) | 28 (80.00) |

p<0.01

### SUPPLEMENTAL METHODS

**Variable Selection and Survival Prediction Models**

The survival function for the Cox proportional hazards model is as follows^1^:

$\lambda\left( t|x_{i} \right)=\lambda_{0}(t)exp\left( \sum_{j} x_{ij}\beta_{j} \right)$ ,	(1.1)

where $\lambda\left( t|x_{i} \right)$ represents the hazard function at time $t$ for the $i$-th patient, $\lambda_{0}(t)$ is the baseline hazard function, $x_{ij}$ represents the $j$-th feature for the $i$-th individual being studied, $\beta_{j}$ represents the corresponding coefficient, and $exp(x\beta$**)** is a baseline scaling factor. The likelihood of an event being observed for the $i$-th individual at time $T_{i}$ can be written as

$L_{i}(\beta) = \frac{exp\left( x_{i}^{T}\beta\right)}{\sum_{k:T_{k}\geq T_{i}} exp\left( x_{k}^{T}\beta\right)}$,	(1.2)

where the summation in the denominator is over the set of $K$ individuals where the event has not occurred before time $T_{i}$ (including for the $i$-th individual). In practice, we assume that all individuals in our study are statistically independent from each other. This allows us to specify the joint probability of all realized events as the partial likelihood, $L(\beta)=\prod_{i:C_{i}=1} L_{i}(\beta)$ where the occurrence of the event is indicated by the indicator $C_{i}=1$. The corresponding partial log-likelihood can then be written as

$l(\beta) = \sum_{i:C_{i}=1} \left[ x_{i}^{T}\beta-log\sum_{k:T_{k}\geq T_{i}} exp\left( x_{k}^{T}\beta\right) \right]$.	(1.3)

The regression coefficients $\beta$ can be estimated by maximizing the partial likelihood. The LASSO adds a restriction to the Cox proportional hazards function in the following manner such that features with no impact on survival will have their coefficients shrunk to 0:

$\hat{\beta}=\min_{\beta}\left\{ -l(\beta) + \lambda\sum_{j} |\beta_{j}| \right\}$ ,	(2.1)

where $l(\beta)$ denotes the partial log-likelihood from Eq. (1.2) and $\lambda$ is a user-specified tuning parameter that we obtain through 5-fold cross-validation. In our study, we select features with non-zero regularized coefficient estimates where

$\left| \hat{\beta}_{j} \right|>0 .$	(2.3)

In contrast, the Bayesian Cox regression iterates through several different models with different combinations of features to find the best fitting model. The sampling distribution of the survival data uses the same partial likelihood as specified in Eq. (1.1)^2^: The posterior inclusion probability (PIP) for each $j$-th feature is calculated by summing up the posterior probabilities of all models for which it is included as variable^2^. This is computed as the following

$PIP_{j}= \sum_{k:\gamma_{jk}=1} p\left( M_{k}|D \right)$ ,	(3.2)

where $M_{k}$ denotes the $k$-th model, $\gamma_{jk}$ is an indicator variable which denotes that the $j$-th feature is included in the $k$-th model, and $D$ is the survival data. Alternatively, the PIP can be written as

$PIP_{j}= p\left( \beta_{j}\neq0 | D \right)$ .	(3.3)
